# Radiotherapy triggers pro-angiogenic signaling in human lung

**DOI:** 10.1101/2024.10.11.617840

**Authors:** Juliette Soulier, Sandra Curras-Alonso, Maxime Dubail, Hugo Laporte, Ayan Mallick, Chloé Lafouasse, Delphine Colin, Jean-François Côté, Jérôme Didier, Christelle Pouliquen, Abdelali Benali, Marco Alifano, Catherine Durdux, Diane Damotte, Marine Lefèvre, Mylène Bohec, Kim Cao, Gilles Créhange, Pierre Verrelle, Nicolas Girard, Agathe Seguin-Givelet, Arturo Londoño-Vallejo, Charles Fouillade

## Abstract

Radiotherapy is one of the main therapeutic options for the treatment of lung cancer. Although highly efficient, radiation cause severe damages to normal tissue and radio-induced toxicities vary from mild pneumonitis to pulmonary fibrosis. The mechanism leading to these toxicities remain unclear. To investigate the molecular responses of human lung to radiotherapy, we analyzed, by single cell RNAseq, lung tissue resected in the vicinity of the tumor (i.e. treated with radiation) and compared the transcriptional profiles of the distinct lung populations from the same patient removed at distance from the tumor (i.e. non-treated with radiation).

Analysis of six lung samples from patients suffering from Pancoast tumor, a rare lung malignancy that requires neo-adjuvant radiotherapy before surgery, revealed a strong induction of VEGF signaling after radiotherapy. Expression of VEGFA, one of the canonical pro-angiogenic ligands, was found upregulated in multiple cell populations in lung exposed to high doses of radiation. Irradiated capillaries, particularly gCap cells, expressing KDR/VEGFR2, present transcriptional profile similar to tip cells, characterized by sprouting and motility capacities. In addition, we identified a sub-population of alveolar macrophages expressing FLT1/VEGFR1, a receptor for VEGFA, in lung tissues treated by radiotherapy. Cell-Cell communication analysis revealed that FLT1/VEGFR1 positive macrophages interact with tip cells after radiotherapy through IL1B-IL1R signaling. Lastly, analysis of mouse single cell dataset confirmed the increase in the proportion of gCap cells presenting a tip-like phenotype after radiation injury.

Altogether, this study describes, at the single cell level, the pro-angiogenic responses of human lung after radiotherapy. These results will lead to a better understanding of the physiopathology of lung radiation injury and may pave the way to optimize treatments to improve patients’ quality of life.

## Introduction

Lung cancer is the leading cause of cancer-related death worldwide[1]. Most of the patients treated for this disease undergo radiation therapy. However, the lung is a sensitive organ to radiation[2], therefore the treatment is often limited by dose of irradiation that the lung can sustain. Radiation of healthy lungs induces damage, radio-induced lung injury (RILI), involving DNA damages[2] and oxidative stress[3], leading to inflammation and processes of wound healing in the lung tissues[4]. In some patients, this early toxicity can evolve into a chronic condition called radio-induced pulmonary fibrosis (RIPF)[5]. RIPF can develop after a RILI through the replacement of normal tissue by scar due to excessive deposition of extracellular matrix, proliferation of fibroblasts [6], disruption of the alveolar structure and vascular damages[7]. These changes prevent gas exchange and lung function, leading to respiratory failure and death[5]. There is for the moment no efficient treatment to cure or even stabilize RIPF[8]. Even if some of the molecular and cellular events that occur in RILI and that lead to RILF have been described[9], the detailed mechanisms and processes leading to fibrosis is not fully understood. Furthermore, most of what is known about RIPF comes from a model of total thoracic irradiation in the mouse[10], mostly because access to fresh human irradiated lung samples remain complex and challenging. In mouse models, it has been described that in reaction to irradiation, healthy tissue goes through processes of epithelial to mesenchymal transition, a switch of profile of the macrophages to either a pro-fibrotic or pro-inflammatory phenotype depending on the population, a transdifferentiation of the AT2 cells to AT1 cells and an increased extracellular matrix production by the myofibroblasts[11].

However, it is important to study the human lung response to irradiation in complement of these studies. Therefore, our goal in this study is to gain a better understanding of the cellular and molecular mechanisms of the human non tumoral lung response to irradiation. In that purpose, we collected samples from patients suffering from Pancoast tumor and who underwent radiotherapy previously to lobectomy[12]. Pancoast tumors are located at the apex of the lung and often invade the sternum [12, 13]. Due to this particularities, a combination of Platinum salts and radiotherapy is used to shrink the tumor prior to surgery[14]. Classically performed after surgery and chemotherapy, Pancoast tumors represent one of the rare cases of treatment with neoadjuvant radiotherapy, allowing us to access previously irradiated human lung. To perform this study, we used single cell RNA sequencing to analyze the molecular responses of human lung to radiotherapy and gain a better understanding of the cellular and molecular mechanisms involved.

After radiation injury, lung vessels are damaged, especially the capillaries. Pro-angiogenic signalling can be involved in the repair processes and angiogenesis. Formation of new vessels requires a complex signalling network involving the vascular endothelial growth factor (VEGF) signaling [15]. At the cellular level, angiogenesis has been shown to include the differentiation of the endothelial cells into two cell states: tip cells and stalk cells. Tip cells express the VEGF receptor KDR (also known as VEGFR2), and sense the VEGF secreted in the environment. Tip cells also develop filipodia, can migrate towards an extracellular gradient of VEGFA, occupying a leading position in angiogenesis. They are followed by the stalk cells that divide to allow elongation of the protruding new vessel [16]. Induction of VEGF signalling in the tip cells triggers the expression of several genes including DLL4. DLL4 ligands then activate NOTCH receptor expressed by the stalk cells. In stalk cells, activation of NOTCH signalling inhibits KDR/VEGFR2 expression and activates FLT1/VEGFR1 expression, maintaining the stalk cell identity [9, 17, 18].

During this study, we will use a single cell RNA sequencing approach to study the cellular and molecular consequences of radiotherapy in human lung obtained from Pancoast patients, particularly the consequence on lung capillaries, a critical population for gas exchanges and lung function.

## Material and methods

### Human samples availability

Freshly resected lung human samples were obtained from six patients undergoing upper lobectomy of a Pancoast tumor who had previously received a neoadjuvant radiotherapy (40- 45 Gy delivered by daily 2 Gy/fraction, considered sufficient to trigger pulmonary fibrosis) concomitant to a chemotherapy with platinum salts on the first and fourth weeks of radiotherapy treatment. Accessibility to the dosimetric computerized tomography (CT)-scans of the patients allowed us to determine a region in the lobe far away from the tumor that did not receive any radiation (NI) and a region next to the tumor that received the highest dose of radiation (IR). A sample of 2 cm^3^ from each of these regions was resected and immediately placed in cold 1x phosphate buffered saline and transported on ice directly to the research lab for single cell dissociation procedure (**fig. 1a**). Accessibility to human samples was achieved in collaboration with Institut Mutualiste Montsouris and Cochin Hospital. Informed consent was obtained from each patient before the surgery.

**Figure 1.**
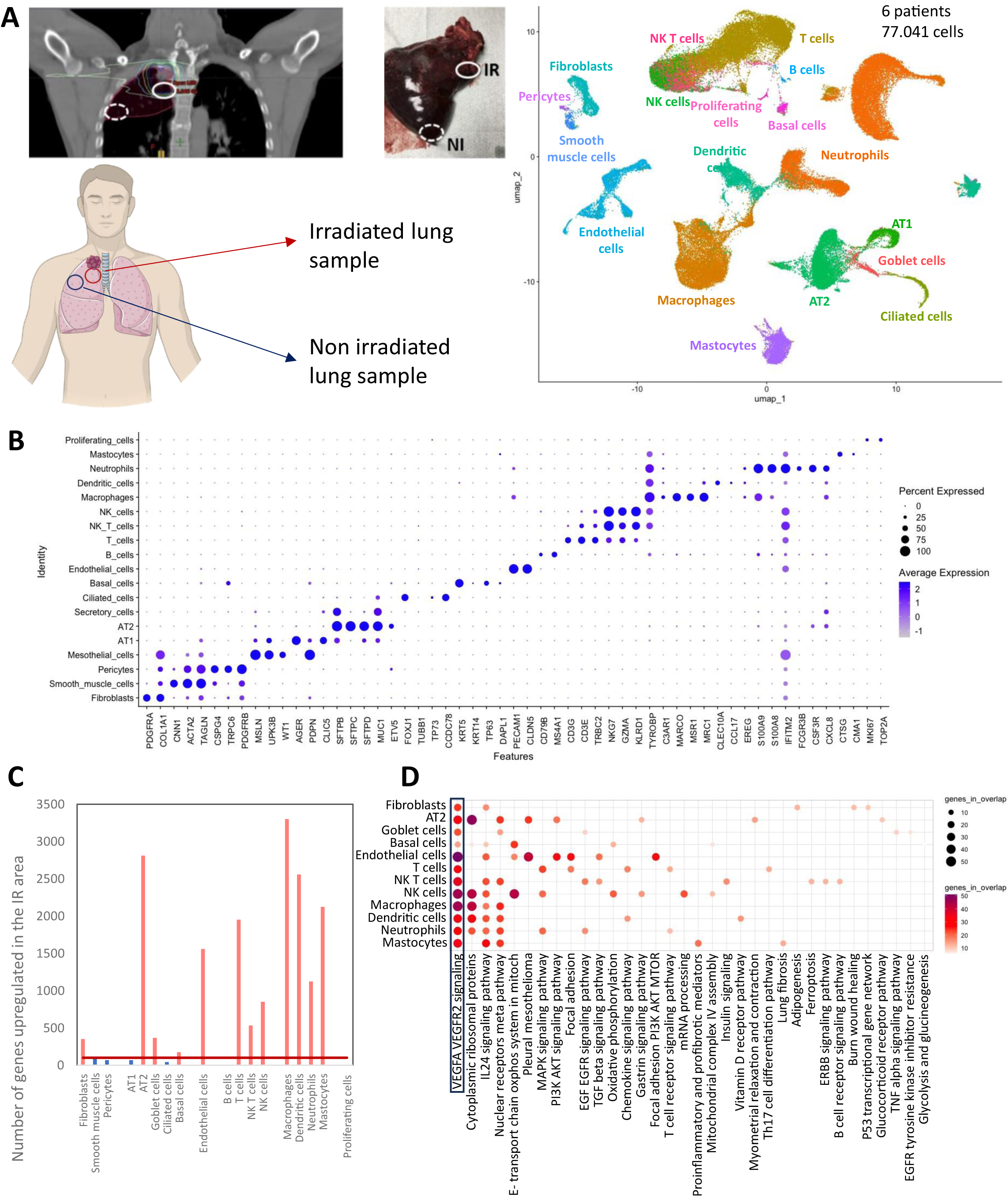
Angiogenesis signaling is upregulated in the irradiated human lung. A: dosimetric CT-scan of a patient who received radiation therapy at Institut Curie and surgery at IMM following a Pancoast tumor, surgical sample (6-8 weeks post radiotherapy), schematic representation of the location of the irradiated and non-irradiated non-tumoral samples from the Pancoast patients and UMAP visualization of 77.048 cells from six patients with 33.328 cells from the NI samples and 43.720 cells from the IR samples, annotated by cell type; B: DotPlot of the expression of characteristic markers of the different lung populations; C: number of genes significantly upregulated in the irradiated area of the lung compared to the irradiated in the different lung populations, the red line is placed a 100 pathways upregulated and the bars in blue represent the cell populations with less than 100 genes upregulated in the irradiated area; D: results of the gene set enrichment analysis of the upregulated genes in the IR area compared to the NI area for the cell populations presenting more than 100 genes upregulated. The pathways are from the WikiPathways database.

### Lung tissue dissociation

Human lung samples were perfused with 1.5 ml of 50 U/ml dispase (Serlabo, WO- LS02100; Sigma Corning, DLW354235) using a 20G needle, followed by 0.5 ml of 1% agarose (Invitrogen, 15510-027) to block the exit of the dispase. Lungs were minced with a scalpel into small pieces and added into 3 ml of 1x DPBS MgCl_2_+ and CaCl_2_+ (Gibco, 14040-091). Then 320 μl of 25 U/ml elastase (Worthington, LS002292) were added and the suspension was homogenized and incubated for 45 min at 37°C with orbital shaking. Enzymatic activity was inhibited with 5 ml of PF10 (1x DPBS containing 10% fetal bovine serum (FBS)) and 20 μl of 0.5 M ethylenediaminetetraacetic acid (EDTA) pH 8 (Invitrogen, AM9260G). Cell suspension was filtered through 100 μm nylon cell strainer (Fisher Scientific, 22363549), which was rinsed with 5 ml of PF10. This was followed by 37.5 μl of 10 mg/ml DNase I (Sigma, D4527-40KU) treatment and incubation on ice for 3 min. Cell suspension was filtered again through a 40 μm nylon cell strainer (Fisher Scientific, 087711) and 5 ml of PF10 were added to rinse it. Samples were centrifuged for 6 min at 150 g and 4°C, pellet was resuspended in red blood cell (RBC) lysis buffer (Roche, 11814389001) and incubated for 90s at room temperature before stopping the lysis with 6 ml of PF10. Then, 500 μl of pure FBS were placed at the bottom of the sample, prior to a final centrifugation for 6 min at 150 g and 4°C. The pellet was resuspended in 1 ml of 1x DPBS containing 0.02% bovine serum albumin (BSA) (Sigma, D4527-40KU) and cell counting was done in a Malassez. Finally, concentration of the samples was adjusted to 1 million cells/ml in 1x DPBS containing 0.02% BSA.

### Droplet based single cell RNA-seq and scRNA seq data analysis

Single cell suspensions were analyzed with the droplet based single cell RNA-seq method proposed by 10x Genomics using the protocol previously described[11]. Raw sequencing data were processed using the CellRanger pipeline (version 3.1.0, 6.0.0 or 7.1.0). Count matrices were analyzed using the Seurat package V5.0.1[19]. For each sample, SoupX[20] was used to remove contamination by ambient RNA and quality controls were performed. The objects from individual patients were annotated using ScArches[21] and the Human Lung Cell Atlas as a reference[22] using a transfer learning method: the ScArches algorithm was trained on the Human Lung Cell Atlas, and the model was then applied to each of our sample individually. Correct annotation of cell populations was verified using the expression of well-known markers. As samples from the different patients presented batch effect when merged, we then integrated the Seurat objects of the different patients using the Seurat method (**fig. 1b**). In order to do that, we first merge all the samples together. The SCTransform function was run on the merged object, with the variables to regress set as the cell cycle score and the percentage of mitochondrial genes. Then the samples from different patients were separated and normalized individually. The 2000 most variable features were calculated and used to set anchors for integration. Finally, the different patient objects were integrated, and the resulting object was scaled.

Different tools and packages were used for the analysis. We used the UMAP method to visualize the data[23].In order to perform the differentially expressed genes analysis (DEG), we used the MAST method[24] to compare two populations, with an adjusted p-value threshold of 0.05, and a logFC threshold of 0.1. Then the upregulated pathways were identified with GSEA and the WikiPathway database[25, 26]. The scoring of “tip” and “stalk” was computed with the AddModuleScore function from Seurat. The comparison between the scores was performed with a wilcox test, with a p-value threshold of 0.05. The intercellular interactions analysis was done with CellPhoneDB V5.0.0[27].

## Results

### The response of human lung to radiation is characterized by activation of pro-angiogenic pathways in several cell compartments

Although most of the dose of irradiation is received by the tumor volume, the surrounding tissue is exposed to doses higher than 45 Gy. Thoracic radiotherapy induces RILI, that impinges cancer treatment possibilities and may evolve towards RIPF, a life-threatening complication of radiotherapy. In order to gain a better understanding of these radio-induced toxicities, we analyzed the response of non-tumoral lung to radiotherapy, by comparing, from the same patient, the transcriptional changes, at the single cell level, from lung tissue resected in in the vicinity of the tumor with healthy tissue sampled from a non-irradiated distal part of the lung (**Fig 1a**). We used single cell RNA sequencing technology to determine the impact of irradiation to the different lungs cell populations. Differential expression analysis between irradiated (IR) and non-irradiated (NI) areas identified genes upregulated in the lung exposed to high doses of radiotherapy (**Fig 1c**), in particular in immune cells (500 to 3.300 overexpressed genes), endothelial cells (EC) (1.500 genes) and type II epithelial (AT2) cells (2.800 genes). Strikingly, the most frequently upregulated pathway across the different cell populations was VEGFA-VEGFR2 signaling, known to be part of the vascular repair/regeneration pathway as well as the classic angiogenesis process[15]. Considering the fact that endothelial cells are a particularly radio-sensitive population[28], it is tempting to speculate that radiation activates the VEGFA-VEGFR2 signaling pathway to enhance vascular repair after radiation injury.

Indeed, blood vessels, in particular capillaries, are crucial for lung function, specifically the gas exchanges that take place in alveoli. To further dissect pro-angiogenic signaling in human lung, we looked in different cell compartments at the level of expression of major ligands (i.e. VEGFA and VEGFB) and their cognate receptors (i.e. FLT1/VEGFR1 and KDR/VEGFR2), in order to identify the sources and targets of this signaling. In the NI area, epithelial cells (mostly AT1), endothelial cells (EC), interstitial macrophages (IM) and mastocytes appeared as the main sources of VEGFA, while alveolar macrophages (AM), smooth muscle cells (SMC) and dendritic cells (DC) appeared to be the main sources of VEGFB (**fig. 2a**). Upon irradiation, the percentages of cells expressing VEGFA gene increased in AT1, IM, fibroblasts and SMC, while for VEGFB, the most important increases were seen in AM, B cells, fibroblasts and AT1 cells (**fig. 2b**). The expression of receptors KDR/VEGFR2 and FLT1/VEGFR1 was largely prevalent in the different types of ECs (**fig. 2c**), and their expression increased upon irradiation, especially in aCap (aerocytes) and gCap (general capillary cells) for KDR/VEGFR2 and in aCap and arterial ECs for FLT1/VEGFR1 (**fig. 2d**). A low percentage of dendritic cells, neutrophils and AM was also expressing FLT1/VEGFR1 and this percentage increased upon irradiation (**fig. 2c-d**). This observation is interesting, since a FLT1+ AM population has been shown to play an important role in some processes during angiogenesis[29]. Together, these results point to a stimulation of the pro-angiogenic signaling that target both lung capillary EC subsets, aCap and gCap. As the gCap have been described as the progenitor population of the endothelial capillary cells compartment[30], we focused subsequent analysis on this cell type.

**Figure 2.**
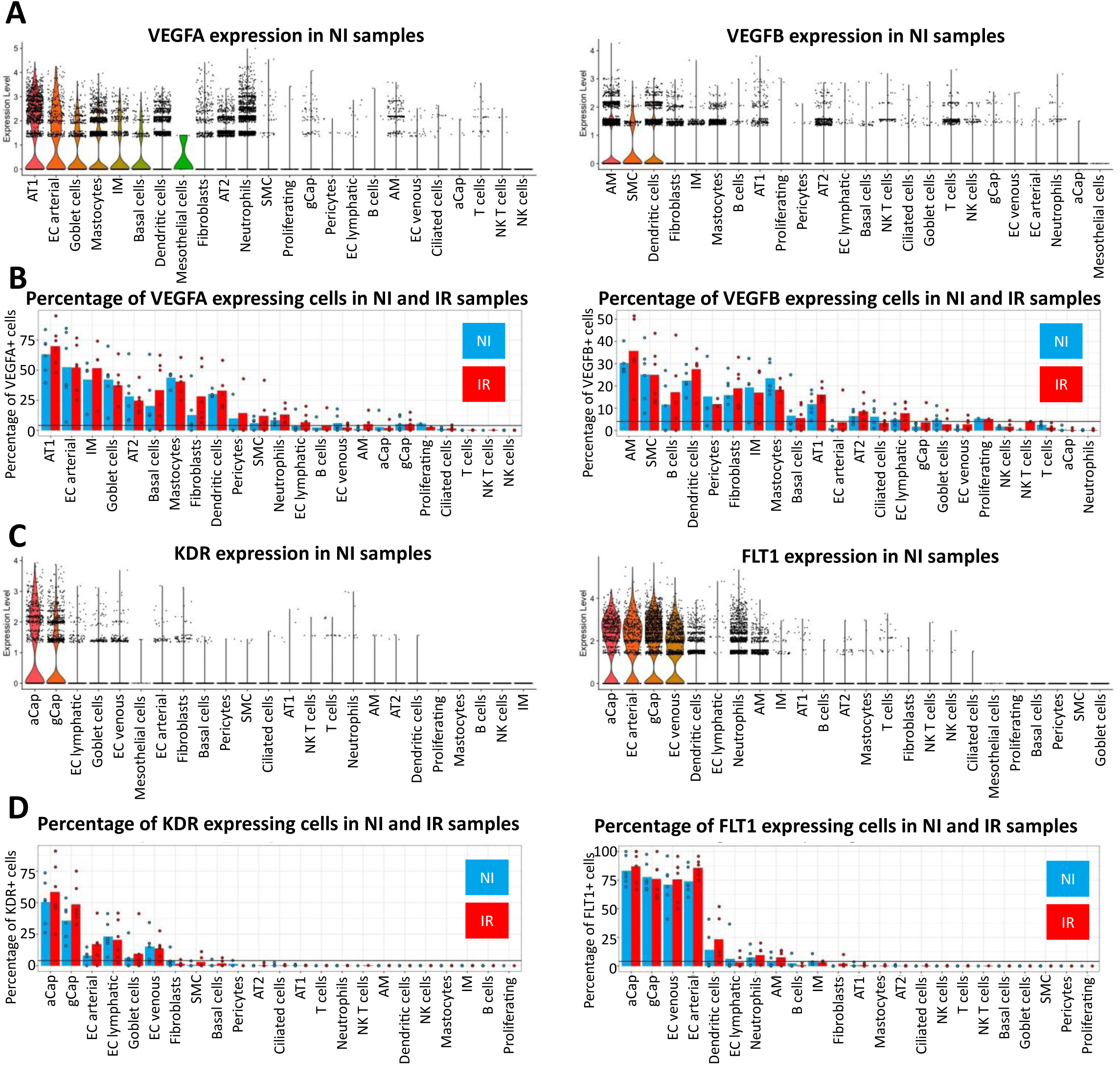
Angiogenic ligands and receptors show an increased expression after irradiation in different cell types. A: violin plots of VEGFA (left panel) and VEGFB (right panel) expression in the non-irradiated area in the different lung populations, sorted by intensity of expression; B: percentage of cells expressing VEGFA (left panel) or VEGFB (right panel) in the irradated or non-irradiated area, with each dot representing a patient; C: violin plots of KDR (left panel) and FLT1 (right panel) expression in the non-irradiated area in the different lung populations, sorted by intensity of expression; D: percentage of cells expressing KDR (left panel) or FLT1 (right panel) in the irradiated or non-irradiated area, with each dot representing a patient.

### Lung gCap ECs display gene expression patterns compatible with a tip phenotype

To better apprehend the consequences of an increase in pro-angiogenic signaling pathway in response to radiotherapy, we further focused our analyses on the different subsets of EC (**Fig. 3a**) (**supplementary fig. 3a**). Since the expressions of either KDR/VEGFR2 or FLT1/VEGFR1 have been shown to be characteristic of two cell states involved in angiogenesis (tip and stalk cells, respectively), we defined two different scores, “tip” and “stalk”, using multiple markers described to be associated with either of these states[31]. The most characteristics marker of the tip cells is Kdr/Vegfr2, and, for the stalk cells, the Flt1/Vegfr1 receptor as well as Ackr1 receptor. However, the distinction of these cell states is subtle and the expression of all markers needs to be taken into account. Based on this analysis, we found a significant increase of the tip score in gCap cells, concomitantly to a significant decrease of the stalk score, in peri-tumoral lung regions exposed to radiotherapy, when compared to non-irradiated areas resected at distance of the Pancoast tumor. These results suggest that irradiated gCap cells respond to the increase in pro-angiogenic signaling by acquiring “tip”-like characteristics. To characterize the transcriptional state of tip cells in the lung after radiotherapy, we compared the transcriptional profiles of gCap cells KDR/VEGFR2 positive versus KDR/VEGFR2 negative. Interestingly, several genes associated with cell motility were upregulated in KDR/VEGFR2 positive tip cells (**fig. 3c**). Furthermore, there is in the IR area a correlation of expression of several motility related genes with the expression of KDR/VEGFR2. For instance, CAV1 promotes EC polarization and movement[32] and ROBO4 is involved in filopodia formation in EC[33] (**supplementary fig. 1a)**. Together, these results support the idea that in response to radiation, lung gCap cells present “tip”-like phenotype that may play an active role in vascular repair process.

**Figure 3.**
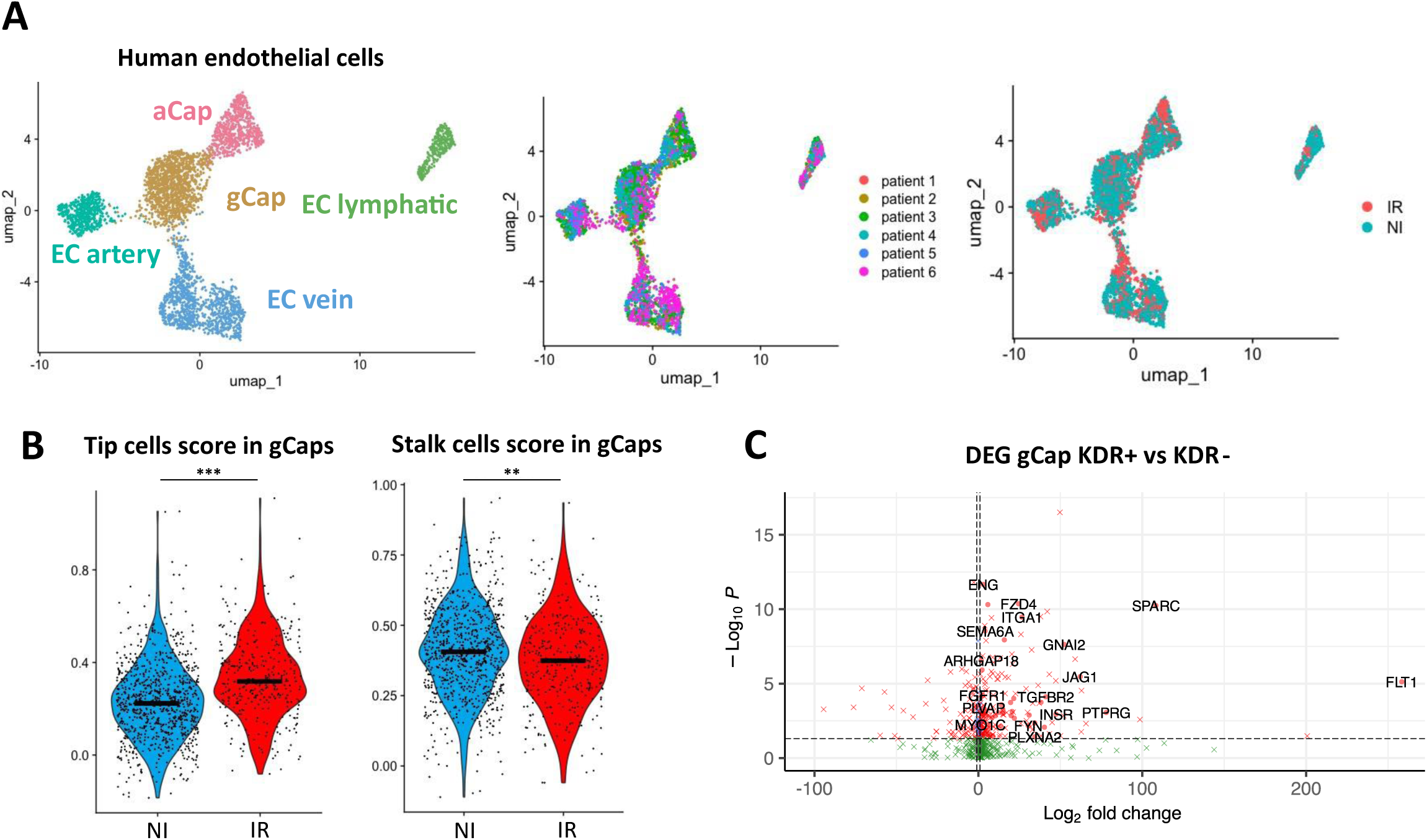
Irradiation induces pro-angiogenesis processes in gCap cells. A: UMAP visualization of 3.648 cells from the different endothelial cell subpopulations annotated by sub cell type, by the patient of origin and by the area (irradiated or non-irradiated) of origin; B: violin plot of the stalk and tip cells scores calculated with the AddModuleScore function, based on the list of markers for these two cell states[31], the black line represents the median; C: volcano plot representation of the DEG analysis of gCap KDR positive versus the gCap KDR negative cells (i.e. genes overexpressed in the gCap KDR positive have a positive fold change). The genes named, and the genes identified with a dot are the genes related to cell motility (from the GOPB cell motility human gene list).

### Alveolar macrophages (AM) present a pro-angiogenic signature in response to radiation

When looking at the expression of angiogenic factors in the different cell populations, we identified a FLT1-positive population amongst resident macrophages. FLT1-positive macrophages have been shown to be crucial for efficient angiogenesis in the lungs[29]. It has been shown that the expression of FLT1/VEGFR1 by circulating monocytes is sufficient to attract them to sites of VEGFA expression and thus stimulate their migration in hypoxic tissue[29, 34]. Therefore, we investigated the presence of this population in human lungs after radiotherapy. Analysis of the dataset revealed an increase of FLT1/VEGFR1 expression specifically in alveolar macrophages (AM) in response to radiation (**fig. 4a-b**). Because FLT1- positive alveolar macrophages have been described to be recruited from circulating monocytes, we next examined the expression level of ITGAM, a specific marker of recruited AM [35]. This marker is expressed by most of the FLT1-positive AM macrophages (**supplementary fig. 2a**), suggesting that radiotherapy enhances the recruitment of monocytes-derived alveolar macrophages from the circulation. Since such macrophages are thought to interact with ECs and support angiogenesis [29, 36], we perform a cell-cell interaction analysis to identify potential signals received by gCap cells presenting a tip-like phenotype. This analysis revealed a potential IL-1b-mediated interaction between the FLT1-positive alveolar macrophages and gCap cells. Indeed, we observed a significant increase of IL-1b expression after radiotherapy amongst the FLT1-positive AM, but not in the FLT1-negative AM (**fig. 4c**).

**Figure 4.**
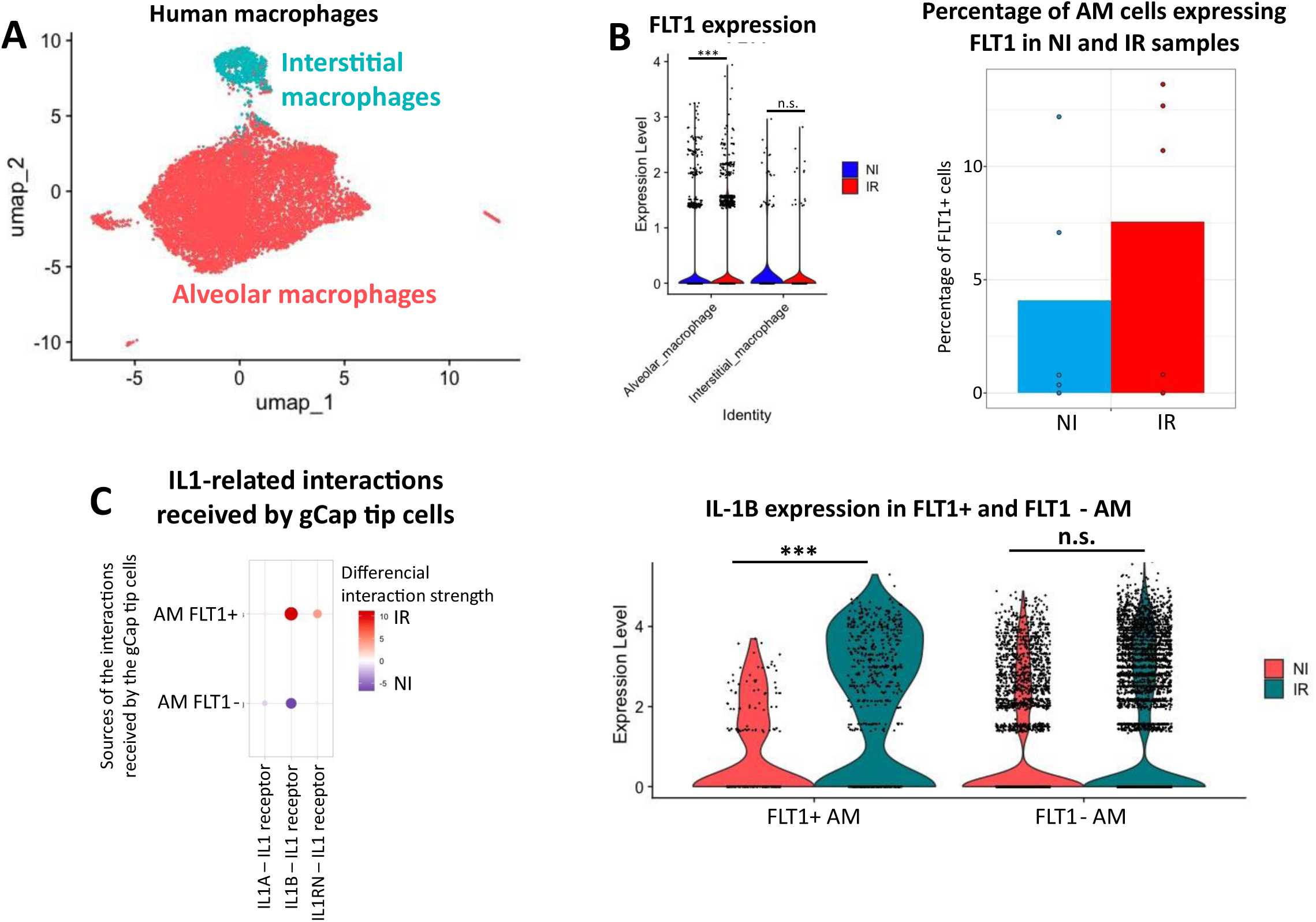
Alveolar macrophages receive and send VEGF signaling after irradiation. A: UMAP visualization of 14.168 cells from the different macrophage subpopulations annotated by sub cell type (4.159 cells from the NI area and 7.009 from the IR area); B: violin plots of FLT1 expression in the non-irradiated and irradiated area in the interstitial and alveolar macrophages and percentage of FLT1 positive alveolar macrophages in non-irradiated and irradiated area, each dot representing a patient; C: DotPlot of the IL-1 related intercellular interactions received by the gCap tip cells and sent by the alveolar macrophages FLT1 positive or negative and violin plots of IL-1 alpha expression in the non-irradiated and irradiated area in the FLT1 positive or FLT1 negative alveolar macrophages.

### Sc RNA-seq analyses in a mouse model of RIFP reveal similar pro-angiogenic responses triggered by radiation

Using previously published lung cell atlas after radiation injury [11], we aimed to characterize such vascular phenotypes dynamically in mouse lung. In this dataset, samples were analyzed every month after either a single fibrogenic (17Gy) or non-fibrogenic (10Gy) dose of radiation delivered to the whole thorax of the mice. After subsetting the endothelial cell populations[30] (**fig. 5a**), we identified in the mouse the same subpopulations identified in patients (**supplementary fig. 3**). In addition, in the mouse dataset, a subpopulation, characterized by high Serpine2 expression, was found in lungs with strong fibrosis, 5 months after exposure to a fibrogenic dose of radiation. The proportion of KDR/VEGFR2 positive gCap cells increased from two months after irradiation (**fig. 5b**). Furthermore, this proportion decreased four months post 10Gy irradiation, while it kept increasing until five months post 17Gy irradiation. Concomitantly, irradiation with a dose of 10Gy triggered an increase of the stalk score at three months before returning to baseline at five months post irradiation. On the contrary, after a 17Gy irradiation, the stalk score increased progressively from three to five months post irradiation (**fig. 5c**).

**Figure 5.**
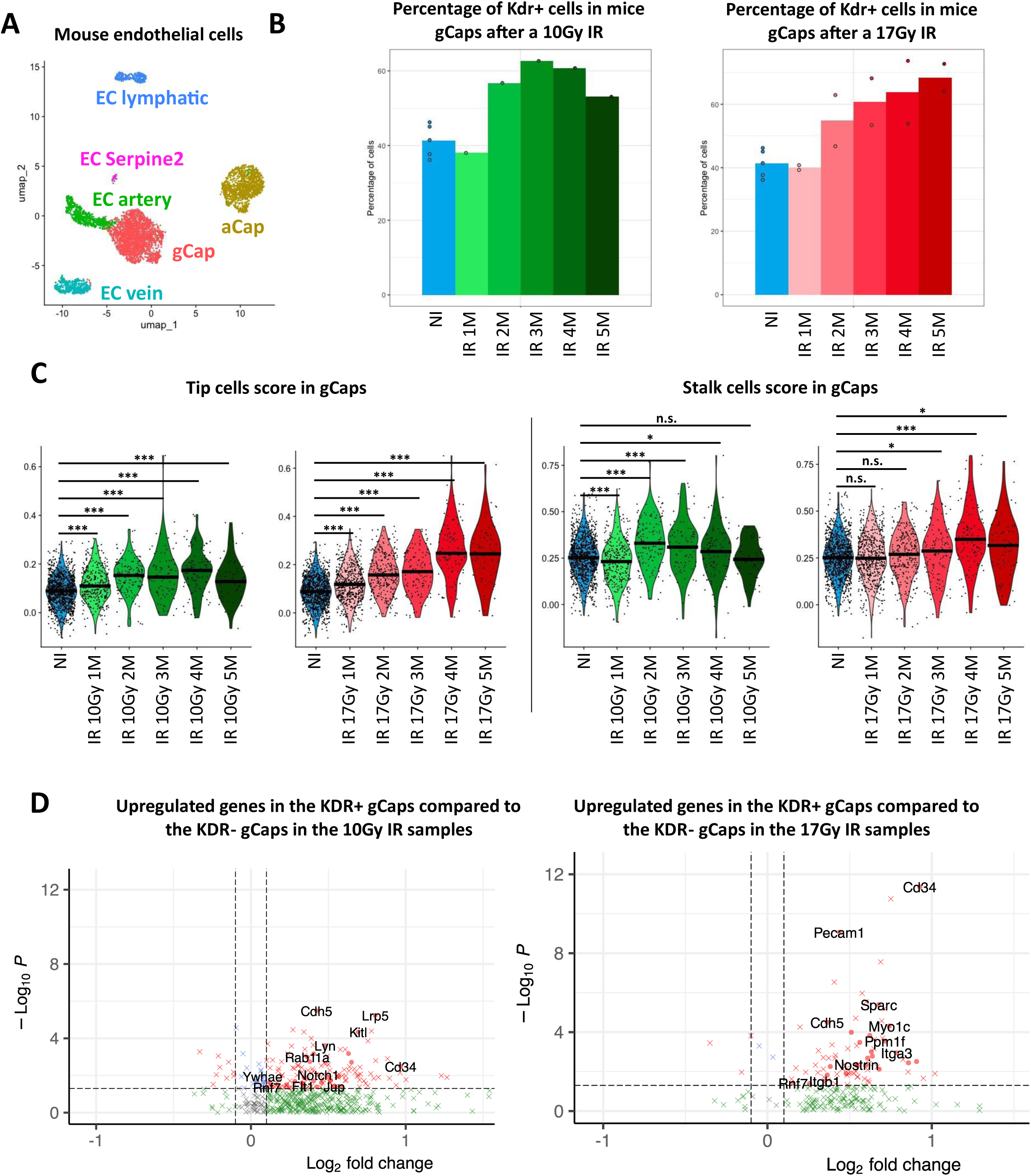
Angiogenesis processes evolve during the months post irradiation in mouse lungs. A: UMAP visualization of 4.958 cells from the different mouse endothelial cell subpopulations annotated by sub cell type; B: proportion KDR+ cells among the gCap cells, in the control mice or after a 10Gy or 17Gy irradiation; C: violin plot of the stalk and tip cells scores calculated based on the list of markers for these two cell states, the black line representing the median; D: volcano plot representation of the DEG analysis of the gCap KDR positive versus the gCap KDR negative cells (i.e. genes overexpressed in the gCap KDR positive cells have a positive fold change) in the 10Gy irradiated gCap cells (left panel) or the 17Gy irradiated gCap cells (right panel). The genes named are the genes related to cell motility (from the GOPB cell motility mouse gene list).

In human, radiotherapy induced the expression of genes related to motility in KDR/VEGFR2 positive gCap cells, in agreement with their tip-like phenotype. To determine if radiation also triggered the expression of motility genes, we analyzed their expression in the mouse lung dataset. The comparison of gCap KDR-positive versus KDR-negative cells from 10Gy irradiated samples showed the upregulation of 31 motility-related genes, while the cells from 17Gy irradiated samples showed only 26 motility-related genes upregulated, like Rab11a [37] or Cd34 [38]. These results confirm that radiation stimulates the expression of motility genes in gCap cells presenting a tip-like phenotype.

## Discussion

Radiotherapy triggers microvasculature damage to the healthy tissues[39], and a functioning vasculature is crucial for the proper function of the lungs. This study aimed at investigating the effects of radiotherapy on human lung vasculature, particularly capillaries, at the single cell level. VEGF signaling is one of the canonical pathway implicated in angiogenesis through VEGFA-KDR interaction[40] and VEGFA-FLT1 and VEGFB-FLT1 interactions[34]. Our single cell RNAseq analysis from lungs resected from patients showed that radiotherapy increased VEGFA/B expression in multiple populations and this pro-angiogenic signal is mainly received by endothelial cells. Furthermore, we identified a subset of lung gCap (i.e. general capillaries) cells that expressed KDR/VEGFR2and present a transcriptional profile similar to the tip cells, described as the leading endothelial phenotype during angiogenic sprouting[31]. In addition, the proportion of alveolar macrophages expressing FLT1 raised after radiotherapy. Cell-Cell Interaction (CCI) analysis predicted this subset of alveolar macrophages interact with the gCap cells, suggesting their implication in the response of lung vasculature to radiotherapy.

After radiotherapy, a high proportion of endothelial cells express FLT1/VEGFR1 but the functions of this receptor are complex. Indeed, FLT1/VEGFR1 has a ten times higher affinity for VEGFA than KDR/VEGFR2, but its kinase activity is ten times lower[41]. Therefore, it has been proposed that FLT1/VEGFR1 acts as a negative regulator of angiogenesis by trapping the VEGFA ligand[34]. However, Flt1/Vegfr1 deficient mice present defects in the sprouting of new vessels [42], indicating that Flt1/Vegfr1 is required for proper angiogenesis. These results suggest that the upregulation of FLT1/VEGFR1 in lung vasculature contributes to angiogenic response that occurs after radiotherapy.

The gCap have been shown to be capillary progenitor cells that are able to regenerate lung capillaries during homeostasis and after injury[30]. Our single cell RNAseq analysis indicated that the proportion of gCap cells expressing KDR/VEGFR2 is increased after radiotherapy. KDR/VEGFR2 has been identified as a marker of tip cells, an endothelial phenotype characterized by its sprouting[31]. Tip cells occupy a leading position in the new vessel formation and are followed by the stalk cells that divide to form the walls of the new vessel[16]. To explore the environment, the tip cells present a high number of filipodia and are able to migrate towards VEGFA gradient. In the tip cells, VEGFA signaling triggers the expression of several genes including DLL4. DLL4 then acts as a ligand for the NOTCH1 receptor expressed by the adjacent endothelial cells. Activation of the Notch pathway in these neighboring cells shifts their transcriptional profile towards a stalk phenotype, activating VEGF receptors expression, forming a VEGF-VEGFR-DLL4-NOTCH-VEGFR feedback loop[17, 31]. In line with this model of angiogenesis, our results suggest that radiotherapy upregulates the expression of VEGFA and that a subset of gCap cells, expressing its cognate receptor (i.e. KDR/VEGFR2), responding to this pro-angiogenic signal, differentiate into tip cells to promote the sprouting of new vessels.

Finally, we identified, in human lung, a population of alveolar macrophages that have been described to be crucial for angiogenesis[29]. After radiotherapy, this particular population may play an important role in the wound healing and repair processes. This subset of macrophages express FLT1/VEGFR1 receptor and are then able to capture VEGFA pro-angiogenic signaling. Interestingly, these macrophages express IL-1, a critical cytokine for angiogenesis. Indeed, it has been shown that supernatant of LPS-treated macrophages is sufficient to stimulate angiogenesis in Matrigel plug injected into mouse interscapular region [43]. The authors also demonstrated that inactivation of IL-1b with IL-1b blocking antibodies is enough to impair this angiogenesis in this model. Altogether, this suggests that radiotherapy increases, in human lung, the proportion of a subset of pro-angiogenic alveolar macrophages FLT1/VEGFR1 positive that secrete IL-1b and contribute to angiogenic responses post-radiotherapy.

Overall, we expect this work will contribute to advance our knowledges of lung responses to radiotherapy, especially its impact on vessels and capillaries. The deciphering of theses mechanisms participates into the effort to gain a comprehensive understanding of the events involved in radiation induced lung injury and could contribute to the discovery of new therapeutic options to fight this side effect of radiotherapy.

---

**Supplementary Figure 1.**
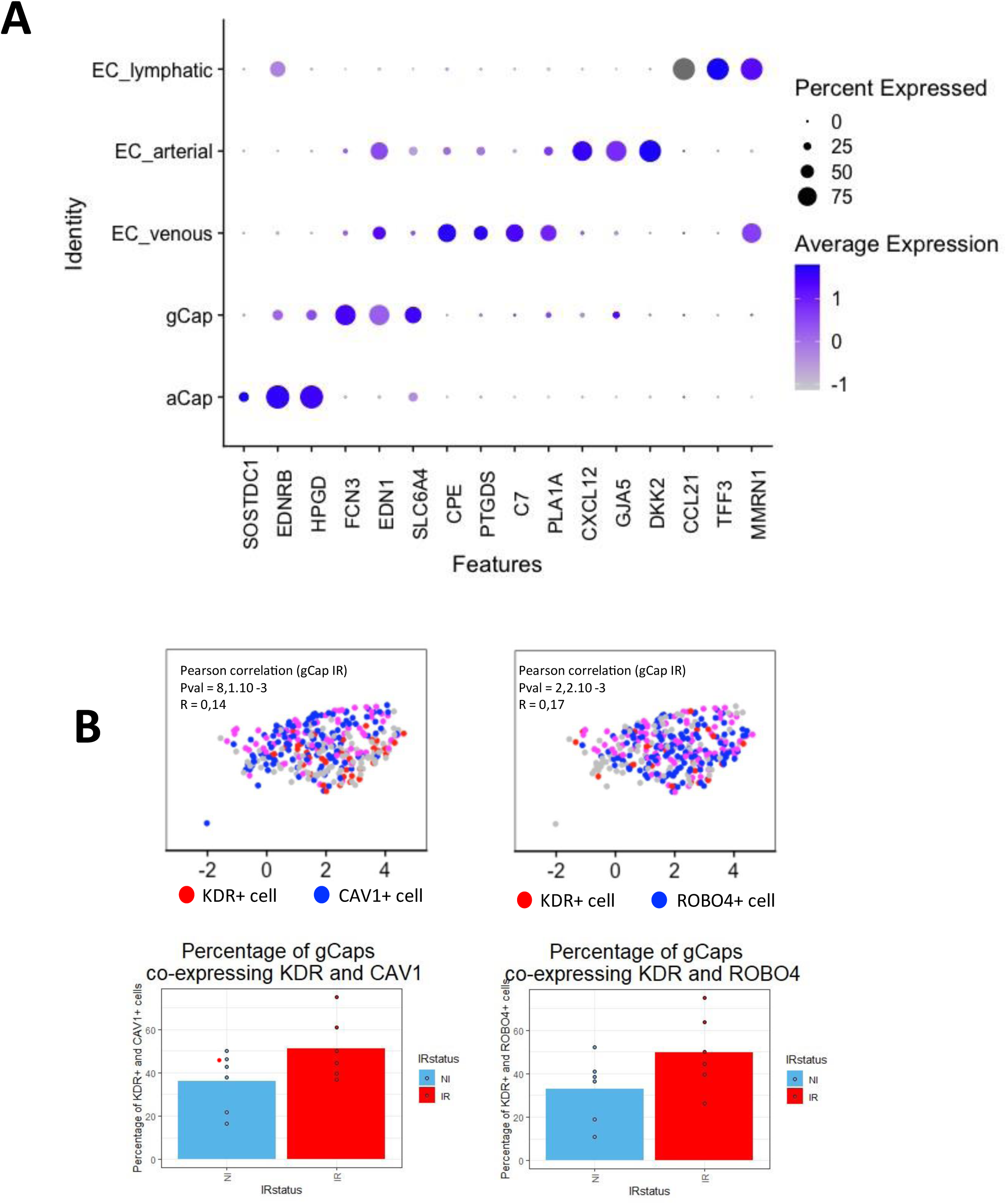
A: DotPlot of the expression of characteristic markers of the different endothelial cell sub populations in human; B: FeaturePlot representing the coexpression of KDR and CAV1 (left panel) or ROBO4 (right panel) in the gCap irradiated cells. Pink cells are coexpressing the genes and percentage of cells co-expressing KDR and CAV1 (left panel) or ROBO4 (right panel) in the non-irradiated or irradiated areas. The correlation and significativity is calculated with a pearson test.

**Supplementary Figure 2.**
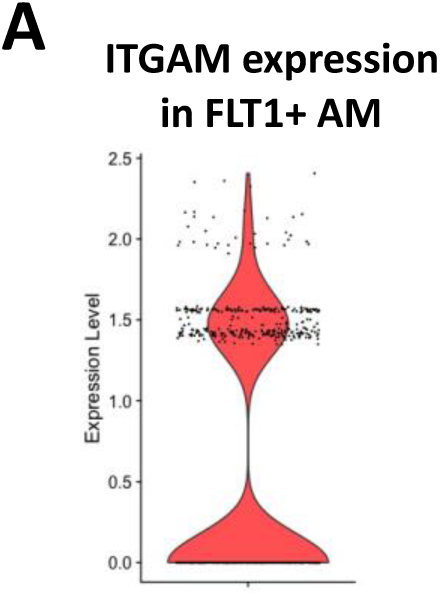
A: violin plot of ITGAM expression in the FLT1 positive alveolar macrophages.

**Supplementary Figure 3.**
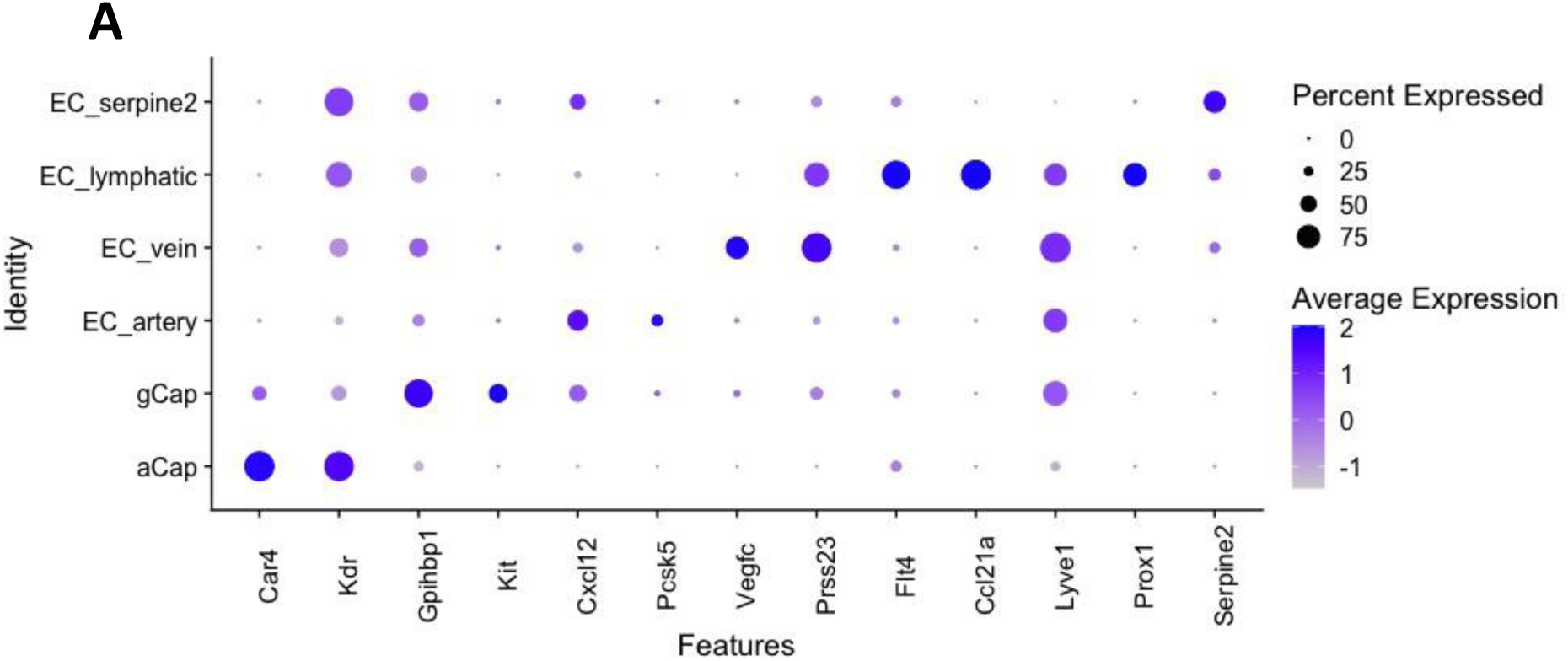
A: DotPlot of the expression of characteristic markers of the different endothelial cell sub populations in mice.

